# Insight into the Impact of Air Flow Rate on Algal-Bacterial Granules: Reactor Performance, Hydrodynamics by Computational Fluid Dynamics (CFD) and Microbial Community Analysis

**DOI:** 10.1101/2024.04.16.589810

**Authors:** Tengge Zhang, Waleed M. M. El-Sayed, Jie Zhang, Leiyu He, Mary Ann Bruns, Meng Wang

## Abstract

Algal-bacterial granules have been drawing attention in wastewater treatment due to their rapid settling ability and efficient nutrient removal performance. This study evaluated the impact of air flow rates on nitrogen removal and the formation of algal-bacterial granules in domestic wastewater treatment. The highest nitrogen removal efficiency was achieved by operating with two separate feedings and the addition of an external carbon source. The higher air flow rate resulted in a higher nitrification rate and produced smaller and more compact granules on average. However, increasing the air flow rate did not necessarily increase extracellular polymeric substances (EPS) production. Computational Fluid Dynamics (CFD) simulations revealed that mechanical mixing was the primary source of shear force. Increasing the air flow rate from 0.2 LPM to 0.5 LPM only yielded a 12% increment in the volume-averaged strain rate. Further analysis of microbial communities showed that changes in bioreactor operation, especially sodium acetate addition and aerations, shifted the microbial community composition. The sodium acetate addition led to the increase of microbial diversity and the relative abundance of denitrifiers such as *Thauera*, while the aeration caused the increasing relative abundances of nitrogen-related genera (such as *Nitrospira*) and the decreasing relative abundances of cyanobacteria and *Chlorella* in the long-term operation of the photobioreactors. Moreover, the decrease in total abundance of grazers and pathogens along with the operation, including *Chytridiomycetes, Sessilida, and Operculariidae*, might result from the shear force and the decrease of prokaryotic species, such as *Chlorella* spp..

**Highlights:**

- A higher air flow rate resulted in a higher nitrification rate.
- Shear stress, microbial composition, and carbon source affected EPS production.
- Increasing the air flow rate from 0.2 to 0.5 LPM led to only 12% of the increment of shear stress.
- Microbial community differed with aeration rate and carbon source.

**Graphical Abstract:** 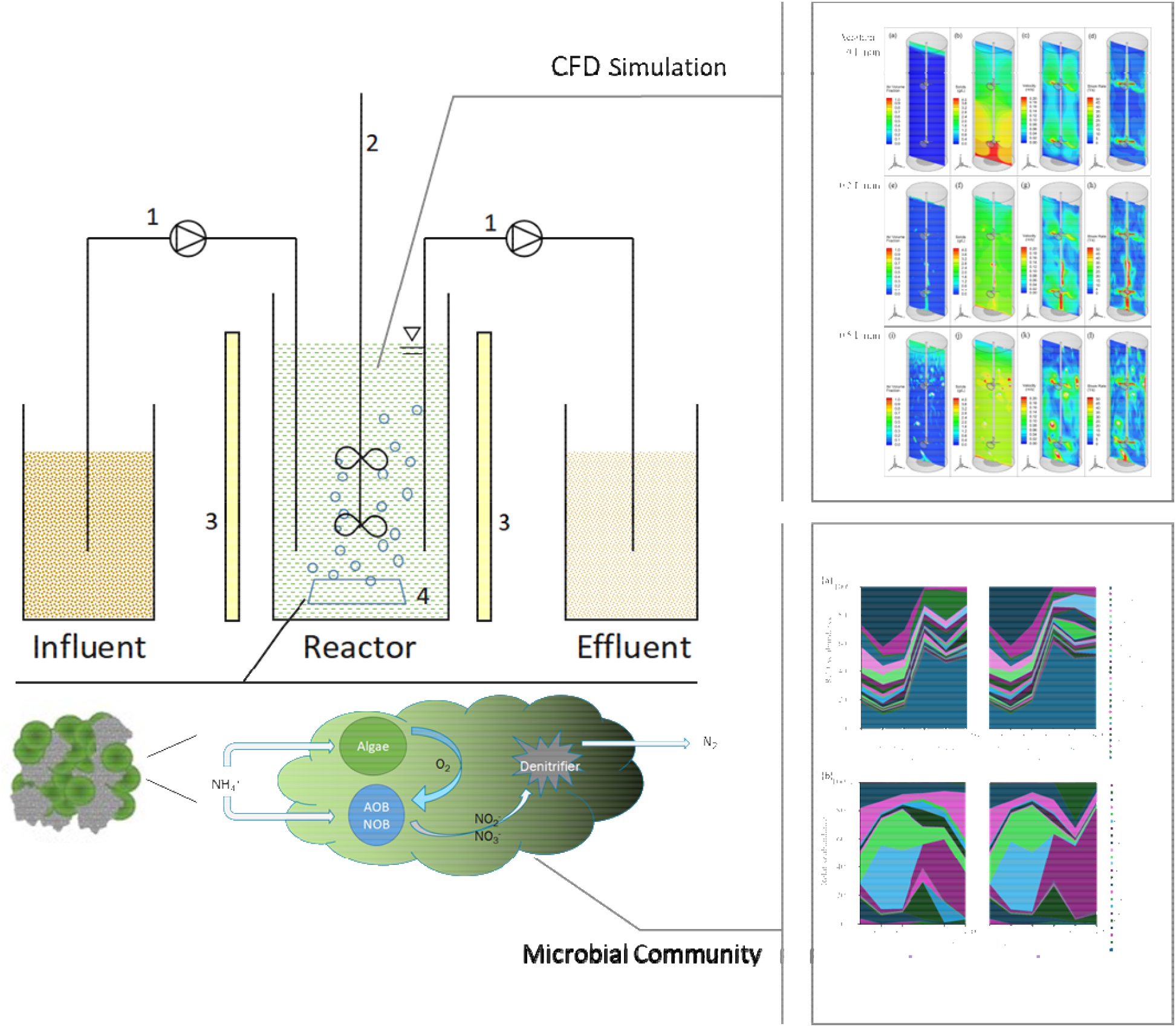

## Introduction

Algal-bacterial consortia have emerged as one of the most promising wastewater treatment technologies for nutrient management (Liu et al. 2017b, Tang et al. 2022). In these systems, oxygen (O_2_) produced by algal photosynthesis can be utilized by nitrifiers and aerobic heterotrophs for ammonium and organic matter removal, respectively. Simultaneously, carbon dioxide (CO_2_) generated by the bio-oxidation of organic matter serves as the carbon source for algal growth. The harvested biomass can be converted to value-added products such as biofuel, bioplastic, and biofertilizer (Arora et al. 2021, Devi et al. 2023). However, biomass harvesting hindered the large-scale application of algae-based systems due to the poor settling ability of algae (Liu and Hong 2021). Algal granulation presents a viable solution to overcome these challenges, offering rapid settling ability, high nutrient removal rate, and low waste of biomass (Uma et al. 2023). Granulation of algal-bacterial consortia can be achieved using either a single microalgal species (such as *Chlorella vulgaris*) or the natural illumination of activated sludge (Petrini et al. 2020). Various factors, including hydraulic selection pressure, bulk substrate concentration, shear force, light intensity, and microbial community composition, have been shown to affect the granulation process (Nuramkhaan et al. 2019, Zhang et al. 2022).

Aeration plays a vital role in the algal-bacterial granulation process to maintain the microbial structure and the metabolism pathways such as nitrification, denitrification, and photosynthesis. High air velocity creates effective whirlpools to enhance granular permeability, leading to the formation of more compact granules (Khan et al. 2013). Prior research showed that high air velocity can provide shear force to stimulate the production of extracellular polymeric substances (EPS), which are critical for the formation and maintenance of granules by influencing cell surface hydrophobicity, cell surface charge, and bridging functions (Nuramkhaan et al. 2019, Wilén et al. 2018). Moreover, operation parameters such as substrate composition, hydraulic retention time (HRT), and hydrodynamic shear force can shift the abundance of EPS-producing microorganisms and re-regulate EPS-production pathways (Liu et al. 2004). Besides aeration, shear force is influenced by water inflow and outflow patterns, mechanical mixer types, and other hydraulic conditions (Tay et al. 2001). These factors contribute to the complexity of shear force analysis and present challenges in the physical measurement of shear force. Computational Fluid Dynamics (CFD) is an advanced simulation tool that has been widely used in many engineering disciplines to reveal the flow field, including the spatial distribution of shear force within fluid systems. Recent studies employing CFD to examine granular sludge reactors (Pan et al 2016; Fan et al 2018; Bastiani et al 2023) affirm its efficacy in exploring the sludge granulation process. Although various studies have discussed the effect of shear force on biomass granulation and reactor performance, there is a lack of quantification of shear force distribution and systematic analysis of its impact on both granular properties and nutrient removal performance in the photobioreactors.

Shear force and other operational conditions can impact microbial community compositions, which in turn affect reactor performance, biomass granulation process, and granular characteristics. It was reported that cyanobacteria played an important role in the formation and maintenance of granules (Peng et al. 2017), which can be enriched under stressful shear force. Species such as *Acinetobacter* spp. and *Thauera* spp. have been shown to increase the removal efficiencies of organic matter and total nitrogen (TN) in wastewater treatment (Ren et al. 2021). Furthermore, the growth of *Chlorophyceae* under hydraulic shear force can also benefit the formation of granules (He et al. 2018). Conversely, zooplankton grazers and predators have been reported to negatively affect the algal-bacterial systems (Molina-Grima et al. 2022, Saleem et al. 2013). However, there remains a gap in the comprehensive understanding of the impacts of operational strategies and air flow rates on the dynamics of microbial communities during the granulation process in the long-term operation of photobioreactors.

This study aims to evaluate the impact of operation conditions, especially the air flow rate, on algal-bacterial granules for domestic wastewater treatment. The specific objectives are to 1) investigate the impact of air flow rate on the performance of photo-sequencing batch reactors (PSBRs); 2) reveal the hydrodynamics and solids behavior of algal-bacterial granular systems; and 3) assess the microbial community shaped by operational conditions.

## 2 Material and methods

### 2.1 Reactor setup

The algal inoculum, obtained from the settling tank of a local wastewater treatment plant, was cultivated in BG11 medium under continuous illumination at 120 μmol/m^2^/s for over a year. To maintain the culture, half of the mixed liquor in each flask was replaced with fresh growth medium every two weeks.

Two identical cylindrical PSBRs with a working volume of 2.5 L (height 0.35 m, diameter 0.11 m) were operated at constant room temperature (25°C) (Fig. 1a).

**Figure 1.**
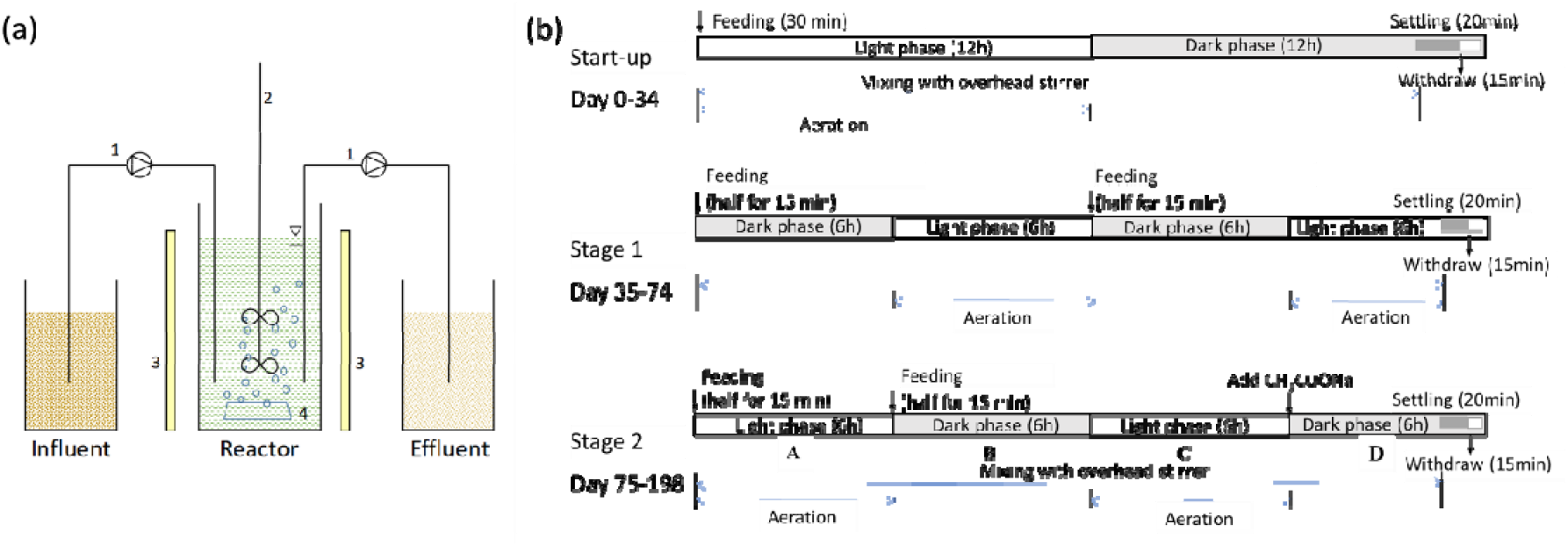
(a) Diagram of the photobioreactors: 1-Pump, 2-Overhead stirrer, 3-Lamp, 4-Aeration. (b) The operation programs of the three stages. Phases A, B, C, and D in stage 2 were denoted for CFD simulation.

Concentrated algal inoculum was seeded to the reactors to achieve a total suspended solid (TSS) concentration of 3 g/L. The photobioreactors were fed with wastewater collected from a local wastewater treatment plant. Overhead mixers were used to keep the biomass in suspension during the reaction stages. Air flow rates of 0.2 liters per minute (LMP) and 0.5 LMP were provided for Reactor 1 (R1-Low) and Reactor 2 (R2-High), respectively. The average light intensity of 200 ± 15 μmol/m^2^/s was provided by LED growth light (BLOOMSPECT, S1000 LED Grow Light), illuminating from the side of the reactors during the light phase. Different durations of light phases and feeding conditions were applied in 3 stages (Fig. 1b). A 12h light/12 h dark cycle was applied during the Start-up stage (Day 0-34). In Stage 1 (Day 35-Day 74), two feeding phases were provided to utilize the organic carbon in the influent to promote denitrification in the second dark phase. In Stage 2 (Day 75-Day 198), an additional external carbon source (sodium acetate) was introduced at the beginning of the second dark phase to promote denitrification. The volumetric exchange ratio was 66% for both reactors. The solids retention time (SRT) was kept at 30 days by decanting a certain volume of mixed liquor before settling.

### 2.2 Analytical methods

The methods for chlorophylls a and b contents, EPS contents, dissolved oxygen (DO), pH, concentrations of NH_4_^+^, NO_2_^-^, and NO_3_^-^, total inorganic nitrogen (TIN), chemical oxygen demand (COD), TSS, volatile suspended solids (VSS), sludge volume index (SVI), and the granular size distribution were described in SI (Text S1, S2 and S3).

### 2.3 Statistical analysis

Basic statistics including mean, standard deviation, and independent group two-sided t-tests were performed in Microsoft Excel. Differences were considered statistically significant when the p-value was less than 0.05.

### 2.4 Nitrogen mass balance

A time-phased study for the 24-hour operating cycle was performed to calculate the nitrogen mass balance on 59 and 69 days for Stage 1 and 158 and 196 days for Stage 2 when the reactor reached the steady state (i.e. NH_4_^+^ removal efficiency reached > 95%). Nitrogen removal was achieved through nitrification, denitrification and assimilation by algae (Rada-Ariza et al. 2017). Two-step nitrification was considered (Eq. 1 and 2). The overall nitrification is represented by Eq. 3 (Liu and Wang 2012):

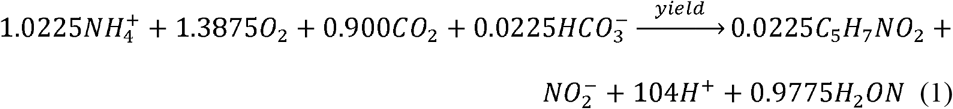

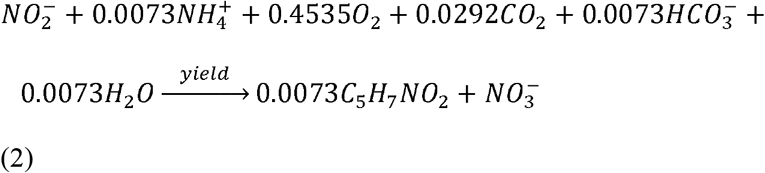

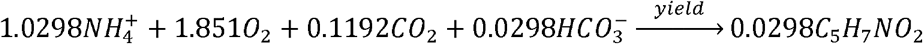

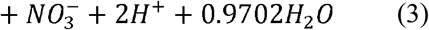

The total mass removal of NH_4_^+^ by nitrification was calculated using Eq. 4, and the total mass of NH_4_^+^ by algae assimilation was calculated by Eq. 5:

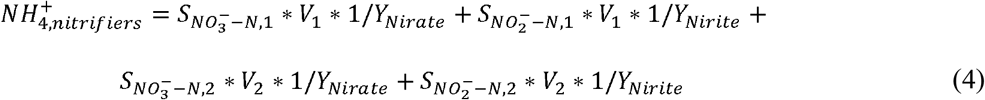

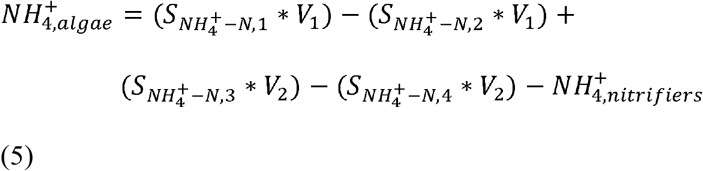

The contributions for NH_4_^+^ removal by nitrifiers and algae were calculated by Eq. 6 and Eq. 7, respectively:

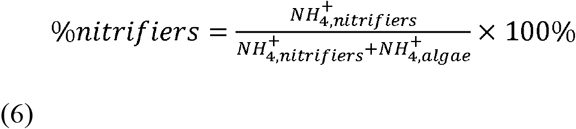

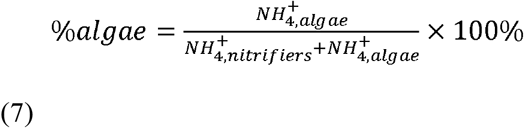

The mass of inorganic nitrogen removed by denitrification (*N*_*denitrification*_) was determined by the amount of NO_x_^-^ removed within an operational cycle. The contribution for TIN removal by denitrification was calculated by Eq. 8:

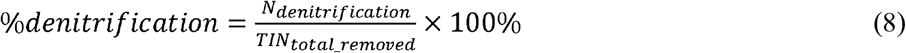

where 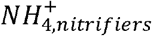 is the total amount of NH_4_^+^ removed by nitrifiers (mg/d), 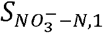 and 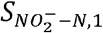 are the NO_3_^-^ and NO_2_^-^ concentration increases from the first feeding to the end of the first light phase (mg/L), *V*_1_ is the liquid volume in the reactor after the first feeding (L), 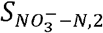 and 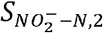 are the NO^-^ and NO_2_^−^ concentration increases from the second feeding to the end of the second light phase (mg/L), *V*_2_ is the liquid volume in the reactor after the second feeding (L), *Y*_*Nitrate*_ is the ratio of nitrate produced through nitrification (0.971 mg NO_3_^-^-N/ mg NH_4_^+^-N based on Eq. 3), *Y*_Nitrite_ is the ratio of nitrite produced through nitritation (0.977 mg NO_2_^-^-N/ mg NH_4_^+^-N based on Eq. 1), 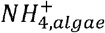 is the total amount of NH_4_^+^ removed by algae (mg/d), 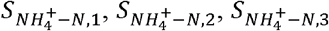, and 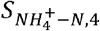 are the NH_4_^+^ concentrations after the first feeding, before the second feeding, after the second feeding, and at the end of the cycle, respectively (mg/L), *TIN*_*total_removed*_ is the total mass of inorganic nitrogen removed during one cycle.

### 2.5 Molecular biology analysis

Seed biomass and biomass samples from each operational stage were collected for 16S and 18S amplicon sequencing. Genomic DNA from each sample was extracted using the FastDNA™ SPIN Kit for Soil following the manufacturer’s protocol. The quantity and quality of extracted DNA samples were assessed using Nanodrop (Thermo Scientific, Massachusetts, USA). Genomic DNA samples were submitted to MR DNA® (Shallowater, TX, USA) for PCR amplification and Next-Generation Sequencing using the Illumina platform. Universal bacterial primers 515/F (5′-GTGCCAGCMGCCGCGGTAA-3′) and 806/R (5′-GGACTACHVGGGTWTCTAAT-3′) were used for 16S rRNA V3-V4 variable region sequencing. The 18S rRNA sequencing primers were EukV4F (5′-CCAGCASCYGCGGTAATTCC-3′) and EukV4R (5′-ACTTTCGTTCTTGATYRA-3′). Denoising of sequences was also performed, and operational taxonomic units (OTUs) were generated. Cut-offs for OTU assignment were defined at a 97% similarity (<3% sequence variation). The data were further processed using methods implemented in QIIME2 (v2022.8) and R (Marizzoni et al. 2020, Xia and Sun 2023). Briefly, sequences were demultiplexed and quality filtered. The demultiplexed and trimmed sequences were denoised with DADA2, a software package that models and corrects Illumina-sequenced amplicon errors within the QIIME2 (v2022.8). OTUs were aligned against the Silva 138 database using the Qiime2-pipeline (Amir et al. 2017). Alpha diversity measures and statistical analyses were performed using R. To identify unclassified OTUs at the genus level, representative sequences from each OTU were blasted against the NCBI’s nucleotide BLAST database for high sequence similarity. This approach allows for the comparison of the unknown sequences to known sequences in the database, helping to assign taxonomic labels to the unclassified OTUs. All sequencing data can be online on NCBI PRJNA1100726.

### 2.6 CFD simulation

#### 2.6.1 Governing Equations and Models

Velocity, air volume fraction, and solids concentration distributions within the reactor can be provided by the CFD simulations. Besides reporting these distributions, the distribution of strain rate is also reported in this study (Fan et al. 2018, Kaya et al. 2023, Pan et al. 2016). Shear stress, arising from the shear force, is the component of the force vector parallel to the material cross-section. Strain rate can be used to represent shear stress since shear stress, τ, can be calculated as

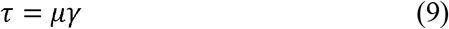

where *γ* is the strain rate, *μ* is the dynamic viscosity of the fluid.

The well-established three-dimensional Volume of Fluid (VOF) transient model was used to simulate the air-liquid two-phase flow in the reactor (van Sint Annaland et al. 2005). The realizable k-ε model was used for modeling the turbulence generated in the reactor. The solids in the reactor were considered passive scalar and solved by a scalar transport equation (Gao 2019):

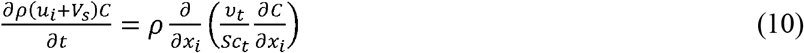

where *C* is the concentration of solids, *v*_*t*_ is eddy viscosity, which is defined as *v*_t_ = *μ*_t_/*ρ*. *v*_*t*_ is computed via the realizable k-ε model. The turbulent Schmidt number, Sct, is taken as 0.7 (Launder, 1978). *V*_s_ is the settling velocity of solids, which is computed by McCorquodale settling model (McCorquodale et al 2005).

The impact of solids on the air-liquid flow was modeled through an extra term added to the momentum equation and turbulence equation. This approach was pioneered by De Clercq (2003) and continues to be used in many studies (Wicklein and Samstag, 2009). Multiple Reference Frame Model (MRF) was used to simulate the rotating stirrer in the reactor. MRF is a steady-state approximation in which individual cell zones can be assigned different speeds. More details about MRF can be found in Ansys Fluent User Guide (2016).

#### 2.6.2 Boundary Conditions and Numerical Tools

Fig. 2 shows an overview of the CFD model of the reactor. The reactors were operated in sequencing batch mode. Two feeding phases in Stages 1 and 2 were introduced in the reactor within 15 mins each and decanting happened in 15 mins. There was no inlet or outlet for water flow during the reaction phases. Thus, snapshot modeling of different reaction phases in one cycle in Stage 2 was performed. Air flowed in through the diffuser at the bottom of the reactor and flowed out through the top of the reactor. Air flow rates at 0, 0.2, and 0.5 LPM were simulated by CFD. Velocity inlet boundary condition was applied to the diffuser for air flow. Pressure outlet boundary condition was applied to the top of the reactor for air venting. No-slip boundary conditions were applied to the walls. The grid used for the simulations has approximately 2 million elements. All CFD simulations were carried out by commercial software ANSYS Fluent (2016). The good modeling practices outlined in Wicklein et al. (2016) were used in this study.

**Figure 2.**
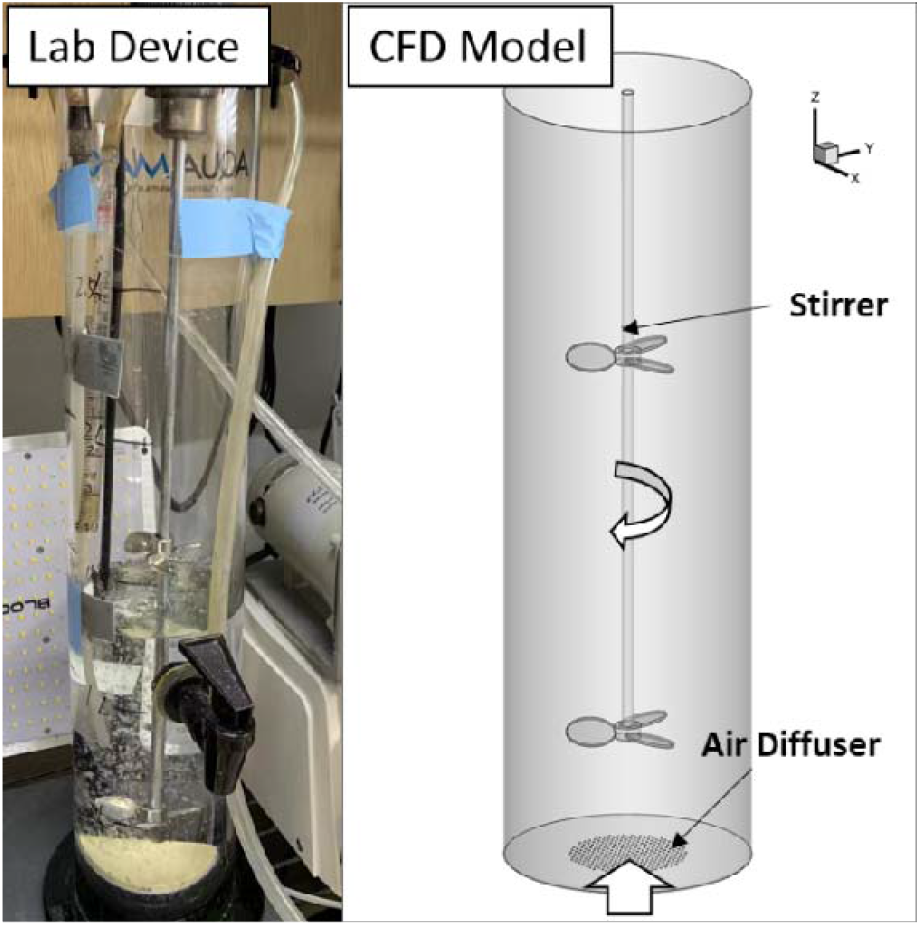
Overview of the lab reactor (left) and CFD model (right).

## 3 Results and discussion

### 3.1 Performance on nutrient removal

During the Start-up stage, both reactors reached over 90% NH_4_^+^-N removal after 20 days (Fig. 3). However, the NO_3_^-^ concentrations of the effluent in the last 10 days were still as high as 37.2±5.0 mg N/L and 39.2±8.6 mg N/L for R1-Low and R2-High, respectively, indicating limited carbon sources for denitrification.

**Figure 3.**
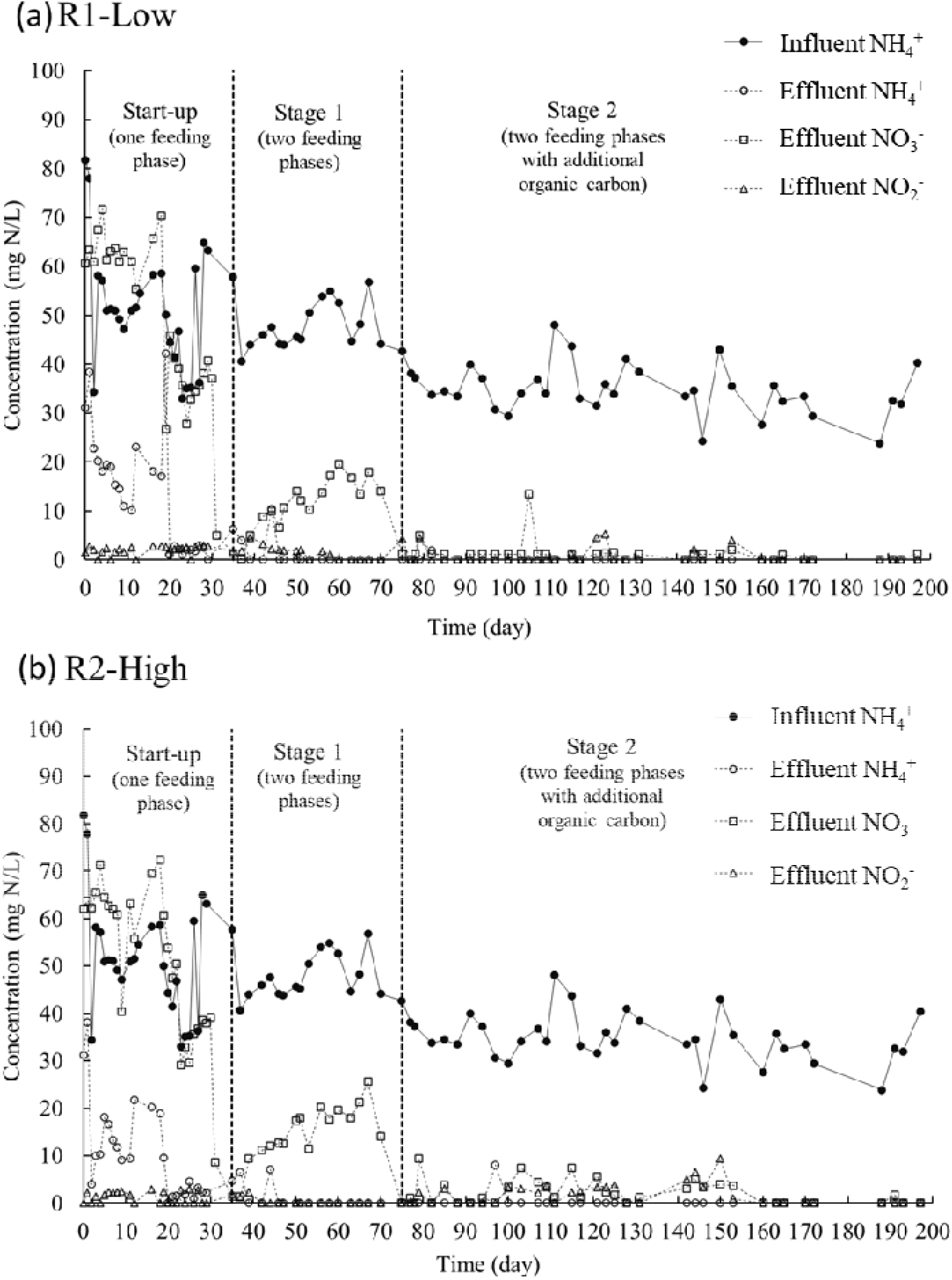
(a) Inorganic N conversion of R1-Low; (b) Inorganic N conversion of R2-High.

The reactors were modified to two-phased feeding after Start-up to utilize the internal carbon source in the influent for denitrification during the dark phases. Results showed that both reactors achieved more than 90% NH_4_^+^-N removal efficiencies (Fig. 3) in Stage 1. Average effluent NO_3_^-^ concentrations were reduced to 11.3±5.6 mg N/L and 14.4±6.5 mg N/L for R1-Low and R2-High respectively, leading to higher TIN removal efficiencies in Stage 1 (more than 68%) than those of Start-up stage. However, the carbon source was still limited for complete nitrogen removal.

An external carbon source (sodium acetate) was added at the beginning of the second dark phase in Stage 2 to promote denitrification. NO_2_^-^-N and NO_3_^-^-N concentrations were not detected in the effluents for R1-Low and R2-High under steady state in Stage 2 (Fig. 3). The addition of external organic carbon source achieved high TIN removal efficiencies of 96% and 90% for R1-Low and R2-High, respectively. Lower average TIN removal efficiency in R2-High was attributed to the insufficient organic carbon source at the beginning of stage 2 caused by the high nitrification rate. Complete denitrification was achieved when an extra carbon source was added from Day 150. Higher aeration led to high shear force and resulted in a smaller size of granules and higher DO concentrations, providing a broader aerobic zone for nitrification (Tomar and Chakraborty 2018), which led to the higher nitrification rate in R2-High.

When using algal-bacterial systems for municipal wastewater treatment, little NO_2_^-^-N and NO_3_^-^-N were removed without organic carbon source addition (Zhou et al. 2022). TIN removal efficiency can reach over 90% in most algal-bacterial granular photobioreactors with sufficient COD for denitrification (Nuramkhaan et al. 2019, Zhou et al. 2022), consistent with the results from this study. In both Stages 1 and 2, NH_4_^+^-N was mainly (61%-82%) removed by nitrification-denitrification process in both reactors, while 18%-39% of NH_4_^+^-N was removed by algal assimilation (Table 1). Similar results were observed from other algal-bacterial granular reactor studies (Liu et al. 2022a, Wang et al. 2023). In this study, R2-High had a higher nitrification rate than R1-Low in both Stages 1 and 2, while R1-Low had a higher percentage of NH_4_^+^-N removed by algal assimilation and a higher Chlorophyll content (15 mg/g VSS and 13 mg/g VSS for Stage 1 and Stage 2 respectively) (Fig. S1). The high algae composition induced high NH_4_^+^-N assimilation.

**Table 1.** Summary of reactor performance.

| Experimental phase | Unit | Reactor | Start-up | Stage 1 | Stage 2 |
| --- | --- | --- | --- | --- | --- |
| Total inorganic nitrogen removal efficiency | % | R1-Low | - | 71.7±8.4 | 96.1±16.8 |
|  |  | R2-High | - | 68.1±8.9 | 89.8±11.2 |
| NH<sub>4</sub><sup>+</sup> removal efficiency | % | R1-Low | 74.7±21.7 | 97.5±6.1 | 99.8±13.2 |
|  |  | R2-High | 81.6±14.5 | 97.7±5.3 | 99.2±4.5 |
| NH<sub>4</sub><sup>+</sup> removed by nitrification* | % | R1-Low | - | 61.2 | 62.0 |
|  |  | R2-High | - | 81.8 | 76.6 |
| NH<sub>4</sub><sup>+</sup> removed by algae* | % | R1-Low | - | 38.8 | 38.0 |
|  |  | R2-High | - | 18.2 | 23.4 |
| Inorganic nitrogen removed by denitrification* | % | R1-Low | - | 30.2 | 53.2 |
|  |  | R2-High | - | 51.7 | 68.2 |
| Nitrification rate* | mg N/L/h | R1-Low | - | 2.9 | 2.2 |
|  |  | R2-High | - | 6.4 | 4.3 |
| COD removal efficiency | % | R1-Low | 49.8±10.1 | 69.0±16.8 | 67.0±20.3 |
|  |  | R2-High | 52.5±4.6 | 69.3±14.8 | 71.6±20.4 |

\*Calculated based on the average values of time-phased data in Fig. S2.

### 3.2 EPS production in algal-bacterial granular reactors

EPS plays an important role in the mass transfer, formation, and stability of granular biomass (Seviour et al. 2009). The average EPS contents of R1-Low were increased from 44±22 mg/g VSS in Stage 1 to 72±12 mg/g VSS in Stage 2. For R2-High, the average EPS contents were 33±1 mg/g VSS in Stage 1, rising to 59±13 mg/g VSS in Stage 2 (Fig. 4). The higher EPS production in Stage 2, especially the increase of tightly bound EPS (TB-EPS), was critical in the granulation process, affecting zeta potential and the hydrophobicity of cells (Wang et al. 2021). The increase in EPS in Stage 2 was probably attributed to the extra sodium acetate addition, which has been shown to be a good organic carbon source for promoting EPS production (Yang et al. 2022). Protein was the dominant component in EPS of the algal-bacterial granules from both reactors. Although the average EPS contents in R1-Low were higher than those of R2-High, no statistical differences were observed. A high air flow rate did not necessarily lead to high EPS production. Other factors such as the microbial community, wastewater composition, and operational conditions also affected EPS production (Liu et al. 2022b).

**Figure 4.**
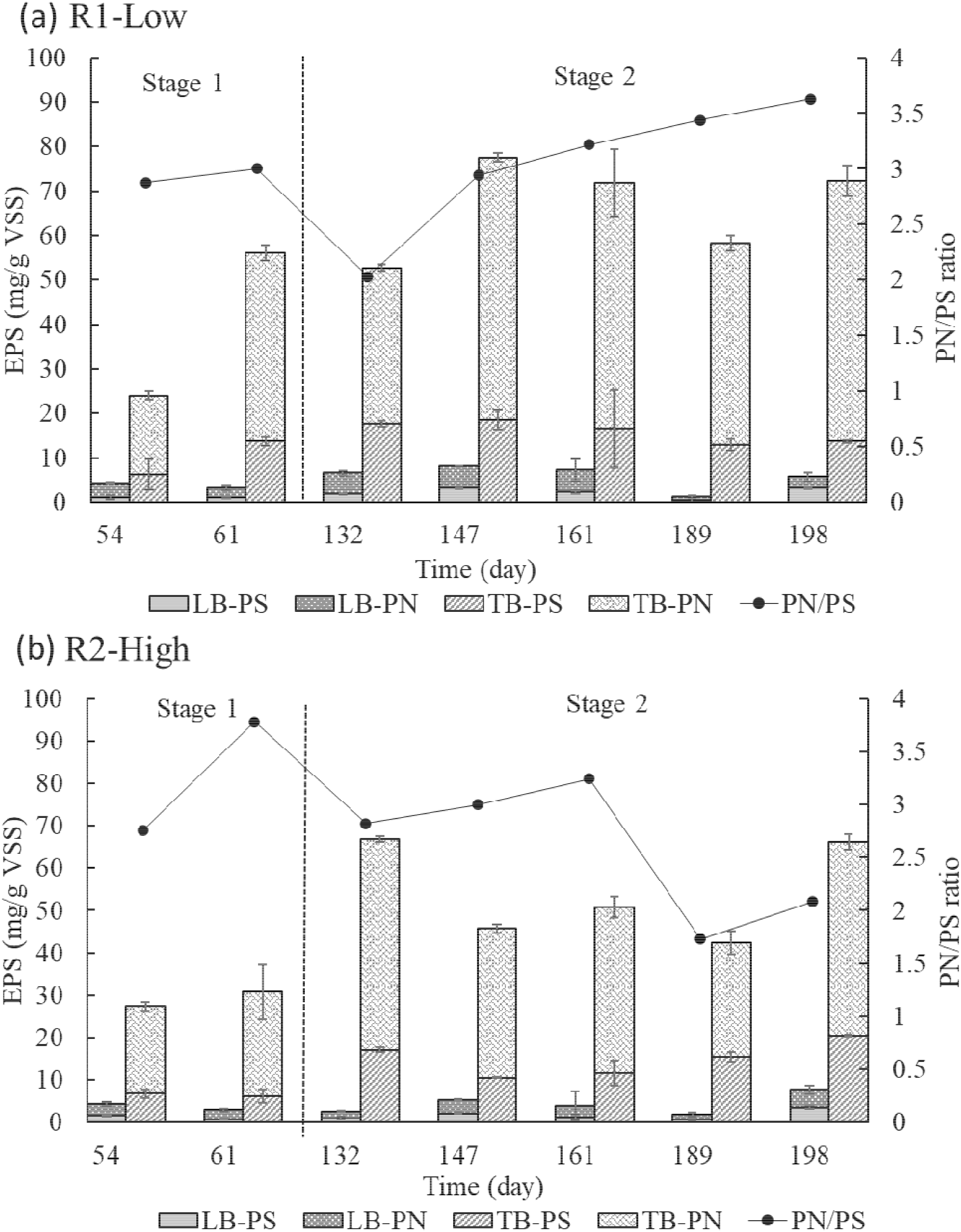
The concentrations of polysaccharides (PS) and protein (PN) in loosely bound EPS (LB-EPS) and tightly bound EPS (TB-EPS), and PN/PS ratio of R1-Low (a) and R2-High (b) in Stage 1 and Stage 2.

The protein (PN)/polysaccharides (PS) ratio of the algal-bacterial granule was reported in the range of 0.5–10 for different influent COD/N ratios (Zhao et al. 2018). Aerobic granular sludge could maintain a good structure at the PN/PS ratio of 4, where the higher protein content contributed to the hydrophobic characteristics of EPS, and polysaccharides helped with the formation of the structure (Liu et al. 2022b). In this study, PN/PS ratios varied from 1.5-4, which are typical values for algal-bacterial granules in maintaining granular structure and stability (Liu et al. 2022b, Wang et al.2020).

### 3.3 Particle size distribution and settleability of the granules

Particle sizes affect nutrient removal performance, biomass settling ability, and biomass recovery efficiency (Tiron et al. 2017). The granules had smaller sizes and better settleability on average in Stage 2 than in Stage 1 for both reactors (Fig. 5). The granules with smaller average size in R2-High also had lower average SVI (Fig. 5). However, there were no significant differences of the size and SVI between the two reactors (p>0.05). In Stage 2, The average sizes were 0.29 mm and 0.25 mm for R1-Low and R2-High, respectively. Similar results were observed in an aerobic granular study with sizes in the range of 0.2-0.6 mm (Chen et al. 2007, Tomar and Chakraborty 2018). The average sizes of R1-Low in both stages were higher than R2-High, which might also contribute to the lower nitrification rate in R1-Low. It was because a lower air flow rate led to larger anoxic/anaerobic regions in the inner granules in R1-Low and less O_2_ transfer from the surface to the core (Kosar et al. 2022). In Stage 2, the sizes of granules in R2-High were smaller than in R1-Low at the beginning and became larger over time. Previous research showed that reactors with higher air flow rate take a longer time (more than 10 days) to form granules, while the granules are more compact and denser than those with lower air flow rate (Tomar and Chakraborty 2018). In this study, elevated air flow rates resulted in higher shear force which will be discussed further in section 3.4. These higher flow rates can enhance the compact granular structure (Chen et al. 2007), causing some loose biomass to detach from the granule surfaces and resulting in a smaller average size at start-up. Along with reactor operations, granules in reactors with higher air flow rate (R2-High) grew larger and the smaller granules with looser structures and lower density were washed out from the reactor with the effluent discharge, resulting in an increase in the average size in this study.

**Figure 5.**
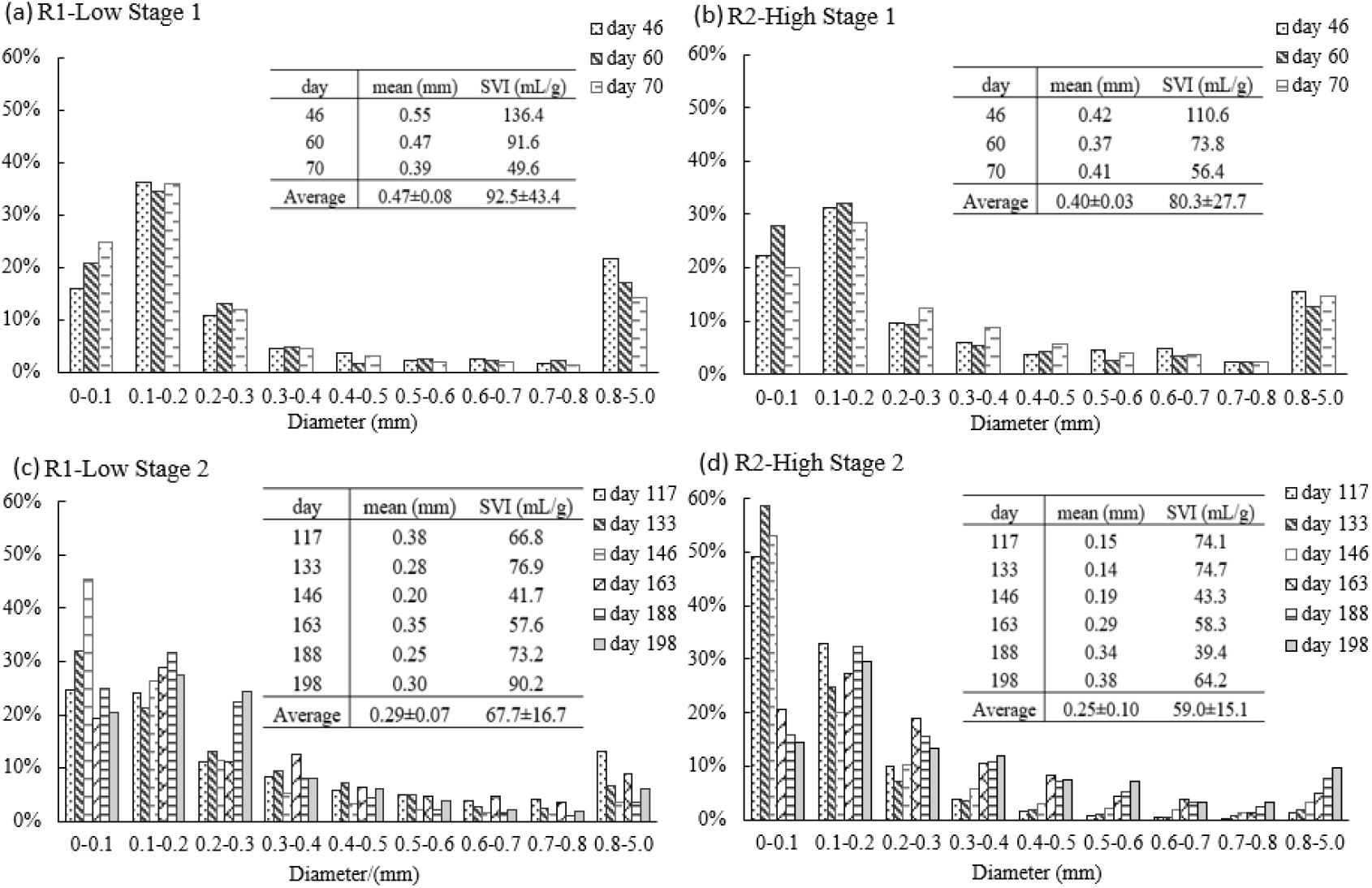
Size distribution and SVI. (a) R1-Low in Stage 1; (b) R1-Low in Stage 2; (c) R2-High in Stage 1; (d) R2-High in Stage 2.

### 3.4 Impact of air flow rate on hydrodynamics and solids distribution in the PSBRs

The primary operation difference between R1-Low and R2-high was the air flow rate. From a physics perspective, the air flow can affect the fluid shear stress in the reactor and affect granular biomass distribution; from a chemistry perspective, air flow can affect the air/oxygen exposure of granules. To analyze the impact of air flow on granular biomass distribution and quantify the contribution of air flow to the total shear stress (note that the mechanical stirrer also contributes to the total shear stress), CFD simulations were conducted for a few scenarios in Stage 2, which had the same operational phases as Stage 1 except for the addition of an external carbon source to promote denitrification. Table 2 shows the operation conditions of the scenarios considered in the CFD simulations, which have similar operation conditions except for the significantly different air flow rates. The CFD analysis is important for system optimization, especially on efficient air utilization and thus high energy efficiency.

**Table 2:** Summary of operation conditions and average strain rates of R1-Low and R2-High in Stages 2B and 2C.

| Reactor | Stage | Rotating Speed (rpm) | Aeration (LPM) | TSS (g/L) | SVI (mL/g) | Volume-Averaged Strain Rate (1/s) |
| --- | --- | --- | --- | --- | --- | --- |
| R1-low | 2B | 130 | 0 | 2.28 | 67.72 | 6.07 |
| R1-low | 2C | 130 | 0.2 | 2.28 | 67.72 | 8.05 |
| R2-high | 2B | 130 | 0 | 2.63 | 59.01 | 5.96 |
| R2-high | 2C | 130 | 0.5 | 2.63 | 59.01 | 9.00 |

#### 3.4.1 Impact of Air Flow on Granular Algae Distribution

Fig. 6a-h compared the distributions of air, granular biomass, velocity, and strain rate at the middle Y-Z plane of R2-High at Stage 2 without and with aeration respectively. Without aeration, a significant stratification of granular biomass concentration was observed (Fig. 6b), that is, a higher granular biomass concentration occurred in the lower region with lower granular biomass concentration in the upper region. In this scenario, the granular biomass distribution was maintained purely by the flow pattern created by the rotating stirrer (Fig. 6c). When aeration was included (Fig. 6e-h), the uplifting air bubbles changed the flow pattern in the reactor and carried solids to the upper region, consequently making granular biomass more uniformly distributed in the reactor. Even at a lower air flow rate, 0.2 LPM, no significant granular biomass stratification was observed (Fig. 6f). Note that granular biomass stratification is not preferable since it can affect light penetration and consequently the growth of algae.

**Figure 6:**
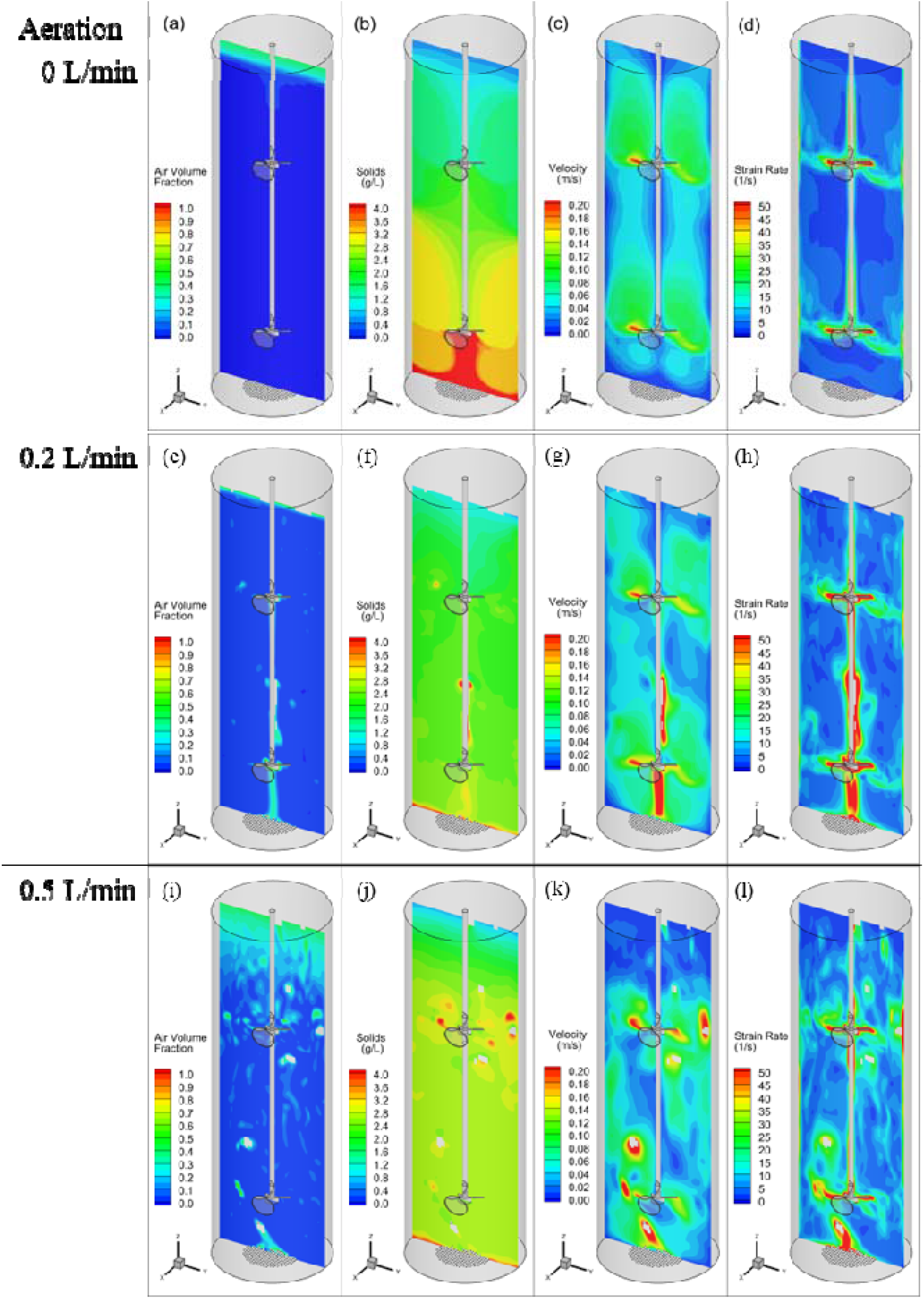
Distributions of air volume fraction, solids concentration, velocity,

#### 3.4.2 Impact of Air Flow on Strain Rate/Shear Stress

The uplifting air bubbles generate additional shear stress, especially in the regions where the stirrer creates downward velocity, like the region below the impellers, as observed in Fig. 6d, 6h, and 6l. The volume-averaged strain rates were calculated and summarized in Table 2. The volume-averaged strain rates versus air flow rates are plotted in Fig. S3. When there was no aeration, the strain rate/shear stress was purely caused by the mechanical stirrer. With air flow rate increasing from 0 to 0.2 LPM, the volume-averaged strain rate increased by 35%. The further increase of air flow rate to 0.5 LPM only brought in another 12% increase in volume-averaged strain rate, indicating further increase of air flow rate may slow the increasing strain rate or shear stress. Overall, in the tests with both stirrer and aeration, the mechanical stirrer provided the majority of shear stress/strain rate (75% at 0.2 LPM and 66 % at 0.5 LPM).

Although the volume-averaged strain rate in R2-high was only about 12% higher than that in R1-low, it caused the mean granule size (0.25 mm) in R2-High to be about 14.8% smaller than that in R1-Low (0.29 mm). This observation is consistent with the previous finding that higher shear stress resulted in smaller granules (Bhatia et al. 2022). Granules of small size tend to have a large total surface area, which promotes the interface mass transfer of gas and substrates, leading to enhanced biomass growth. Moreover, a higher flow rate increases the exposure of biomass to air/oxygen, which may also promote the nitrification process and the growth of nitrifiers. In this case, the impact of shear stress and the growth of biomass shaped the granules simultaneously. As observed in Fig. 5, the mean granular size was consistently lower in R2-high compared to that in R1-low from day 117-163 in Stage 2, indicating fluid shear dominantly impacted the granular size in the early days. However, the trend was reversed in the later stage of the operation (data from days 188 and 198), indicating that the growth of biomass played a vital role in the later stages of granulation, offsetting the impact of shear stress. and strain rate in the reactor under the scenarios listed in Table 2: (a-d) correspond to the No-aeration scenario; (e-h) correspond to the scenario with aeration at 0.2 LPM; (i-l) correspond to the scenario with aeration at 0.5 LPM. (Note that the regions with air volume fraction greater than 0.5 are considered as air phase and blanked in the images).

### 3.5 Microbial community analysis

#### 3.5.1 The distribution of microbial community and its relationship with granular properties

The addition of carbon sources led to higher diversities of bacteria in Stage 2 for both reactors (Table S1). Acetate can be directly converted to acetyl-CoA and then be involved in the tricarboxylic acid cycle (TCA cycle), while some other carbon sources, such as ethanol, are firstly oxidized to acetate and then begin the same biochemical pathway (Sun et al. 2016). It means that acetate can be degraded by denitrifiers more easily than other external carbon sources, promoting metabolism and growth and resulting in higher microbial diversity (de Sousa Rollemberg et al. 2019). The diversity of eukaryotic microorganisms varied (decreased in Stage 1 and increased in Stage 2) for R1-Low. However, the diversity of eukaryotes in R2-High decreased along with the reactor operations in Stage 2, indicating that the combination of the addition of the carbon source and high air flow rates introduced potential stressors for certain algae, resulting in a decrease in diversity.

16S rRNA analysis indicated the cyanobacteria *Nodosilinea* spp. had the highest relative abundance in the seed sludge (Fig. 7). However, their relative abundance decreased precipitously in Stage 2 for both reactors. *Acinetobacter* spp. and *Thauera* spp. became the dominant genera in Stage 2, which might result from the addition of acetate as an extra organic carbon source (de Sousa Rollemberg et al. 2019, Sun et al. 2016). *Acinetobacter* sp. had a synergistic relationship with algae by contributing to the exchange of carbon dioxide and oxygen and improved removal efficiencies of COD and TN in wastewater (Liu et al. 2017a). *Thauera* can perform both denitrification and COD removal (Ren et al. 2021), and support granule formation by producing EPS (Cydzik-Kwiatkowska and Zielińska 2016). The relative abundances of nitrite oxidizing bacteria, *Nitrospira* spp., increased from 0.04% in the seed to 0.2%-1% and 0.6-2% in Stage 2 for R1-Low and R2-High, respectively. It was reported that *Nitrospira* was the dominant genus of nitrite-oxidizing bacteria under high DO conditions, resulting in high NO_2_ ^-^ removal (Fu et al. 2023). Higher abundances of *Nitrospira* spp. contributed to the higher nitrification rate in R2-High.

**Figure 7.**
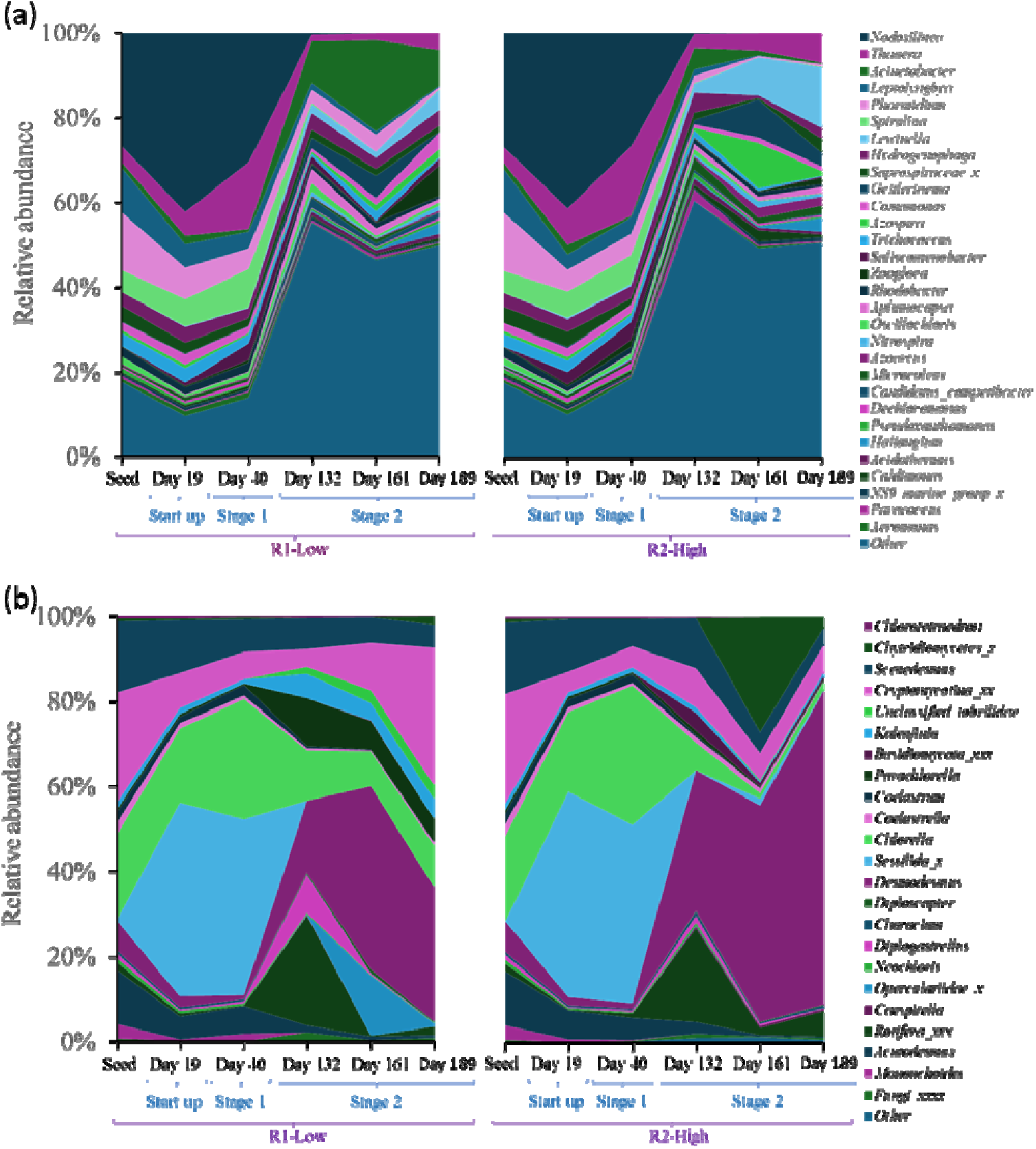
Microbial community composition at the genus level as determined by (a) 16S RNA sequencing of biomass; (b) 18S RNA sequencing of biomass. (Color labels in the figure are arranged from top to bottom)

Cyanobacteria, particularly filamentous *Nodosilinea* spp., *Leptolyngbya* spp., *Phormidium* spp., and *Spirulina* spp. play an important role in granulation by enhancing the formation and stability of granules (Peng et al. 2017), and exhibited a significant presence (56% in total) in the seed. However, cyanobacterial abundance decreased to less than 5% for both reactors in Stage 2. The presence of *Nodosilinea* spp. in the granular sludge promotes the development of granules with increased hydrophobicity surface area, settling velocity, and specific gravity (Shi et al. 2010). In both reactors, the Start-up stage and Stage 1 showed similar abundances of *Nodosilinea* (27-42%). In Stage 2, a notable reduction in the relative abundance of *Nodosilinea* (0.04-0.3%) might contributed to the decrease in granular sizes.

18S rRNA analysis indicated that for the seed, *Scenedesmus, Cryptomycotina, Chlorella*, and *Acutodesmus* were the main eukaryotic genera. The genus *Chlorella* increased from 16%-19% in the Start-up stage to 28% and 33% in Stage 1 for R1-Low and R2-High, respectively. However, *Chlorella* spp. decreased to 8%-12% and 1%-6% in Stage 2 for R1-Low and R2-High, respectively. In Stage 2, *Desmodesmus* spp. increased significantly, and the relative abundances at the end of Stage 2 were 32% and 74% for R1-Low and R2-High, respectively. This was consistent with previous research, where *Desmodesmus* spp. was naturally selected to be the most abundant microorganism in batch reactors for urban wastewater treatment (Samorì et al., 2013).

The correlation analysis between reactor performance and microbial genera revealed distinctive patterns (Fig. S4). *Nodosilinea* spp., *Leptolyngbya* spp., *Telotrochiduim* spp. and *Trebouxiophyceae* spp. showed a positive relationship with particle size and SVI. The genera *Acinetobacter, Anaerolineae, Cephalothrix, Peritrichia*, and *Triplonchida* exhibited a positive relationship with EPS production. Overall, the correlation analysis provides valuable insights into the interconnected dynamics of nutrient removal efficiency, microbial community composition, and operational conditions, shedding light on the complex interactions within the studied consortia.

#### 3.5.2 Dynamics of Zooplankton Grazers and Predators in Algal-Bacterial Photobioreactors

A range of potential zooplankton grazers and predators were identified based on 18S rRNA amplicon sequencing in different stages of both reactors, which have previously been reported to have a negative impact on algal-bacterial systems, causing productivity loss or product contaminant (Molina-Grima et al., 2022). The top zooplankton grazers or predators included members from protozoa (e.g., Operculariidae_x), ciliates (e.g., *Sessilida* spp.), multicellular animals (e.g., rotifers), and fungi including *Diplogastrellus* spp. *Basidiomycota_xxx* spp., *Cryptomycotina_xx* spp. and *Chytridiomycetes_x* spp. (Fig. 8). Overall, *Chytridiomycetes* spp., *Sessilida* spp., *Operculariidae* spp., and protozoa varied with different air flow rates and carbon sources.

**Figure 8.**
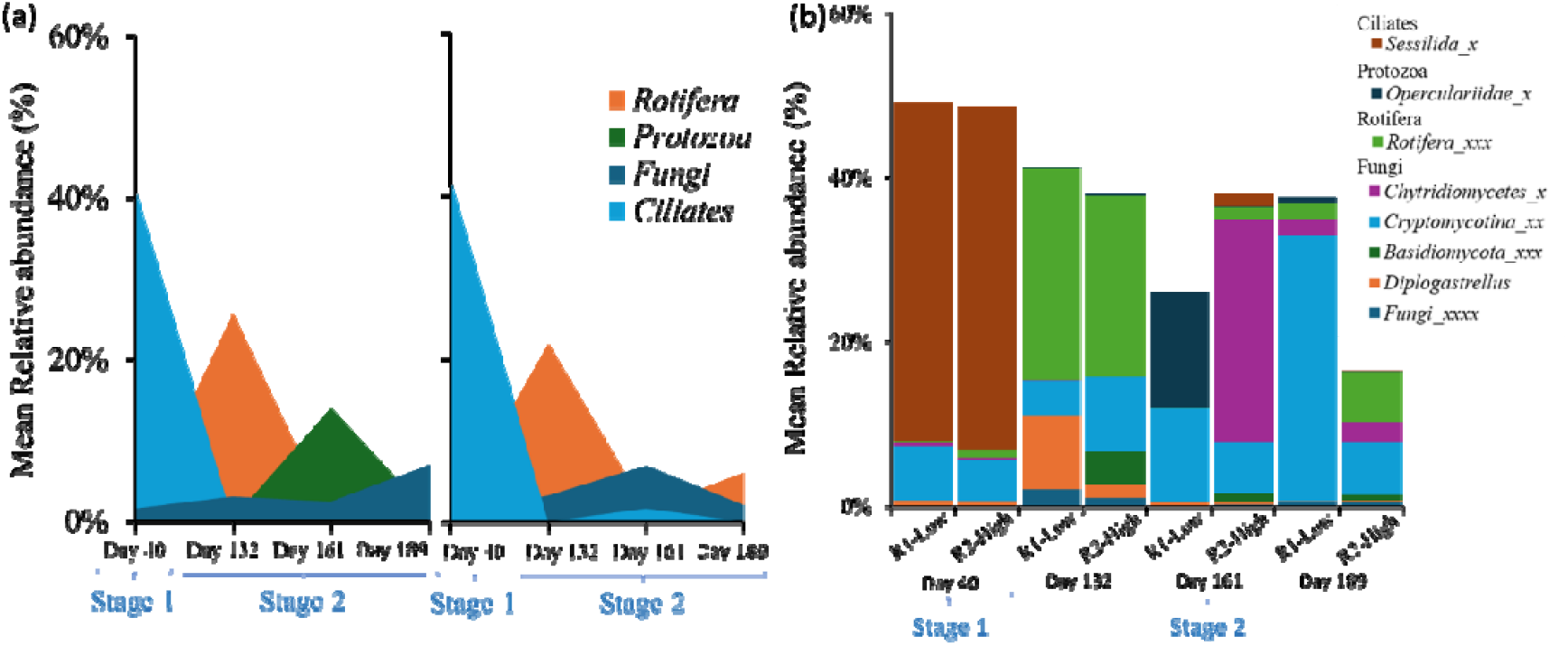
Taxonomic diversity: (a) Groups of potential grazers or predators and fungal pathogens, frequently observed in photobioreactors; (b) Predominant taxonomic levels of grazers or predators.

In Stage 1, ciliates were observed with high main relative abundances of 41% and 42% for R1-Low and R2-High, respectively. In Stage 2, in R1-Low, *Cryptomycotina_xx* spp. (Fungi) showed high mean relative abundance, while other fungal genera, rotifera, and protozoa had lower mean relative abundances. In R2-High, fungi including *Chytridiomycetes* spp. (10%) and *Basidiomycota* spp. (2%) were observed with moderate mean relative abundances. This could be attributed to the decrease of *Chlorella* spp., which could be grazed by *Chytridiomycetes* spp. as a host-specific parasite and among the most pathogenic fungal groups for algal populations (Frenken et al. 2018). In the Start-up stage and Stage 1, *Sessilida* became dominant in both reactors (41%-48%). The mean relative abundance of *Sessilida* reached a minimum at the end of Stage 2 in both reactors (almost 0). This could be explained by an increase in rotifer populations (from 0.2%-2% to 2%-6%), which could be ingesting *Sessilida* individuals (Li et al. 2013). The total relative abundance of grazers or predators decreased significantly in Stage 2, while it was lower in R2-High than R1-Low at the end, which might be because hydrodynamic shear stress, provided by stirrer and aeration, can disrupt zooplankton (Montemezzani et al. 2017).

## 4 Conclusion

The study demonstrated that aeration and operational strategies impacted the nutrient removal performance, the granular properties, and the microbial community of algal-bacterial granular photobioreactors.

- Over 97% of NH_4_^+^ removal was achieved when two feeding phases were applied to treat domestic wastewater.
- The addition of external carbon sources achieved over 90% TN removal.
- Higher air flow rates promoted nitrification by enhanced gas transfer, leading to smaller and more compact granules on average due to increased shear stress.
- CFD simulation indicated that mechanical mixing accounted for most of the shear force, and increasing the air flow rate from 0.2 LPM to 0.5 LPM resulted in only a 12% increase in the volume-averaged strain rate.
- Aeration and additional carbon sources shaped the microbial community, leading to the decrease of cyanobacteria and *chlorella*, and the increase of genera for nitrogen removal and EPS-related microorganisms.
- The decrease in the total abundance of grazers and pathogens might result from the decreased abundance of prokaryotic species and the shear force in the reactors.

The findings suggest that the lower air flow rate is the optimal choice, as it resulted in similar nutrient removal performance and granular properties, while significantly reducing energy costs. Moreover, the study highlights the importance of understanding the interplay between operational conditions, microbial community dynamics, and granular properties to optimize these systems’ performance and stability.

## Acknowledgment

This material is based upon work supported by the Penn State Institute of Energy and the Environment (IEE) seed grant program. Any opinions, findings, conclusions, or recommendations expressed in this material are those of the authors and do not necessarily reflect the views of the sponsors.

