## Appendix A. Supplementary data for "Insight into the Impact of Air Flow Rate on Algal-Bacterial Granules: Reactor Performance, Hydrodynamics by Computational Fluid Dynamics (CFD) and Microbial Community Analysis"

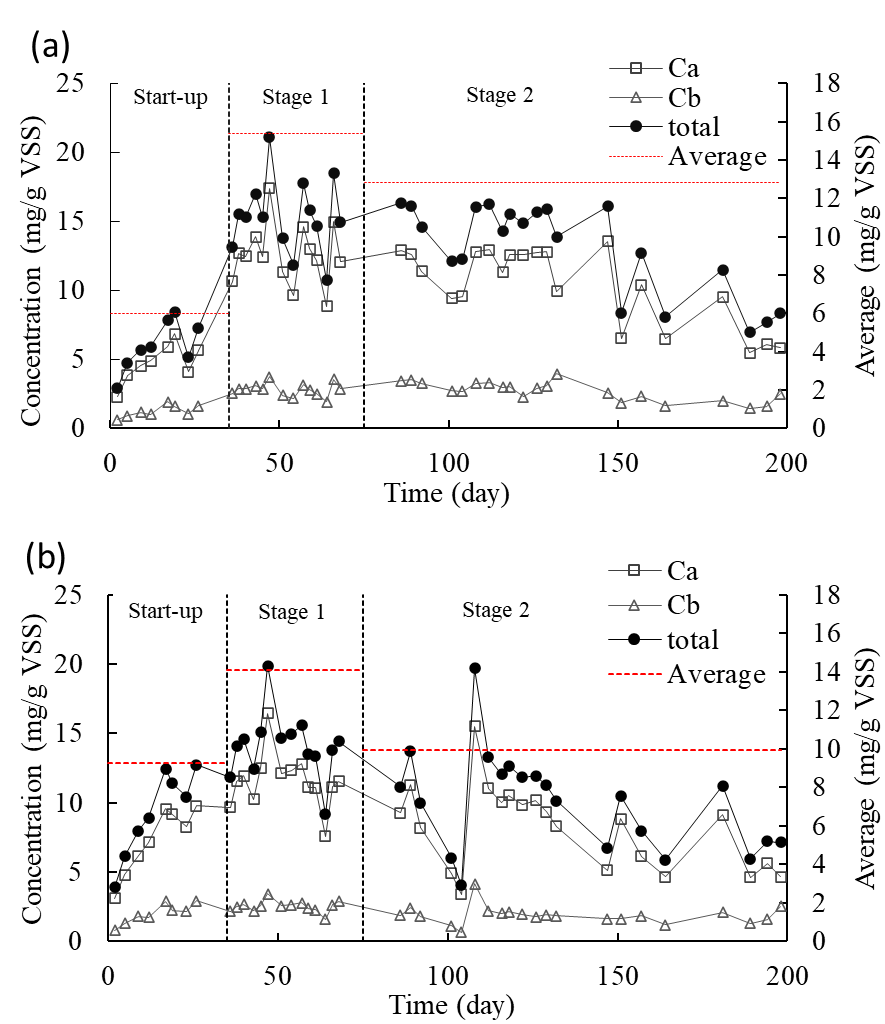

Figure S1. Chlorophyll contents of the biomass for (a) R1-Low and (b) R2-High.

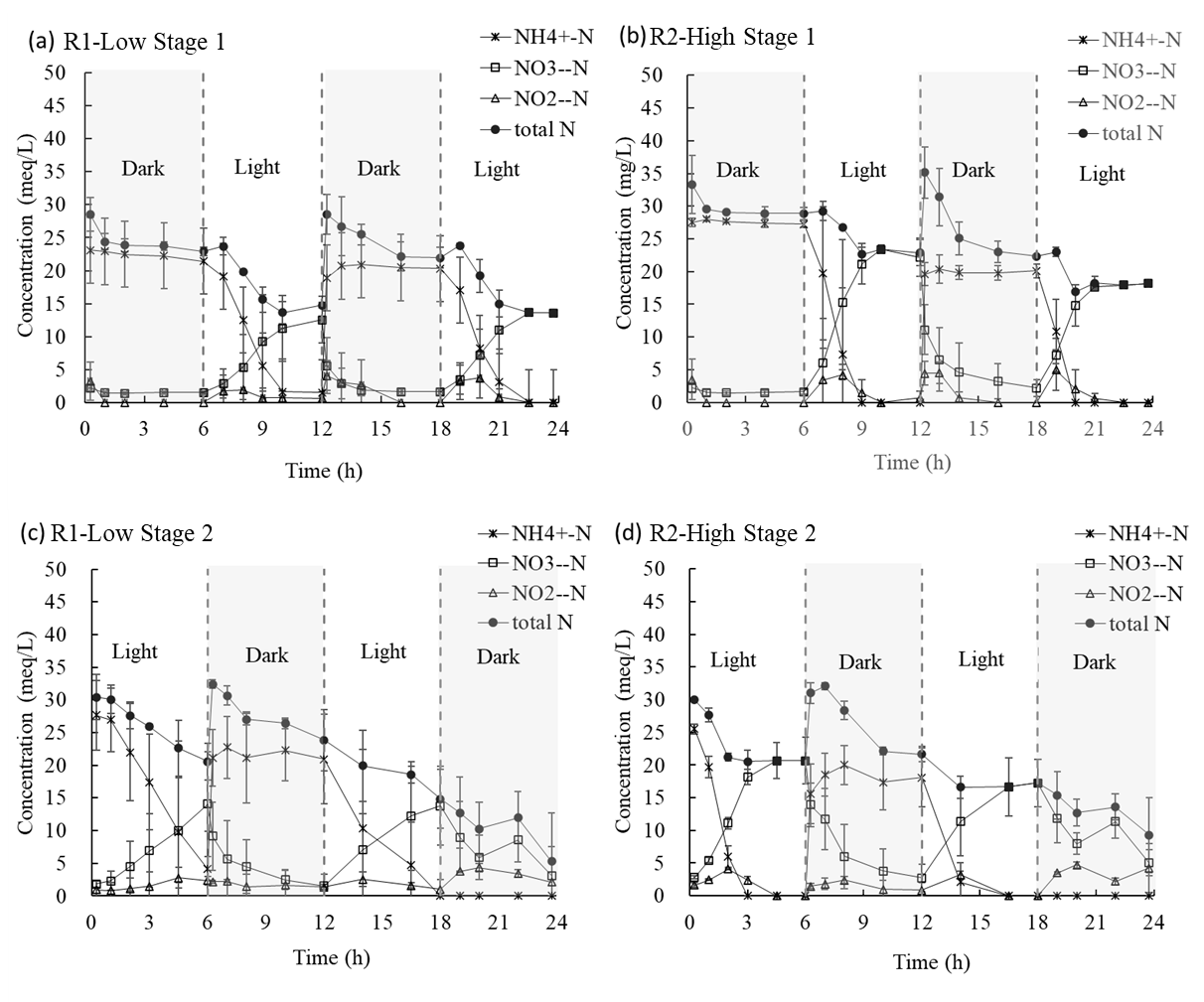

Figure S2. Time-phased inorganic N conversion of both reactors for one cycle in Stage 1 (a&b) and Stage 2 (c&d)

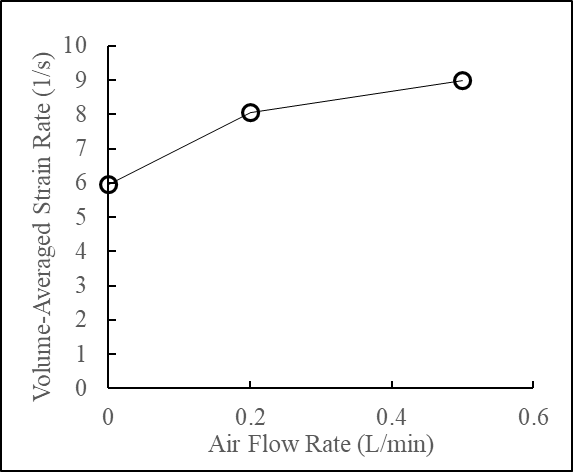

**Figure S3. CFD predicted Volume-averaged Strain Rate versus Air Flow Rate.**

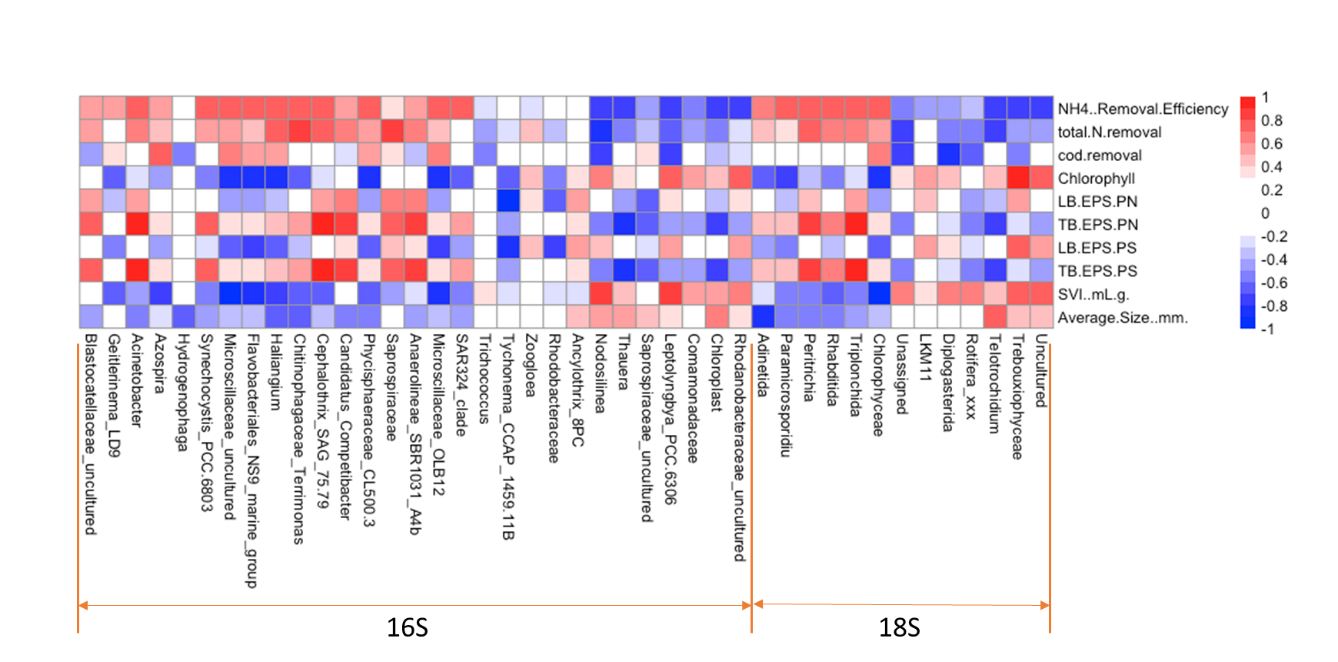

Figure S4. Correlation heatmap analysis between the reactor performance and the predominant microbial at the genus level based on 16S and 18S amplicon sequencing analysis.

Table S1. Alpha diversity values (Shannon and Simpson) for different stages related to 18S Amplicon sequencing.

| RNA analysis | | | 16S rRNA | | 18S rRNA | |
| --- | --- | --- | --- | --- | --- | --- |
| Stage | Day | Reactor | Shannon | Simpson | Shannon | Simpson |
| Seed | 0 | - | 3.845 | 0.910 | 1.894 | 0.832 |
| Start-up | 19 | R1-Low | 3.184 | 0.811 | 1.311 | 0.594 |
|  |  | R2-High | 3.116 | 0.816 | 1.223 | 0.570 |
| Stage 1 | 40 | R1-Low | 3.442 | 0.874 | 1.332 | 0.631 |
|  |  | R2-High | 3.595 | 0.894 | 1.299 | 0.620 |
| Stage 2 | 132 | R1-Low | 4.778 | 0.980 | 1.691 | 0.769 |
|  |  | R2-High | 4.886 | 0.986 | 1.538 | 0.726 |
|  | 161 | R1-Low | 4.415 | 0.957 | 1.512 | 0.684 |
|  |  | R2-High | 4.418 | 0.971 | 1.156 | 0.567 |
|  | 189 | R1-Low | 4.743 | 0.977 | 1.563 | 0.734 |
|  |  | R2-High | 4.465 | 0.964 | 0.716 | 0.294 |

Text S1: Method for chlorophyll analysis

Chlorophylls a and b were measured using a modified spectrophotometric method with alcohol extraction (ISO 1996, Wintermans and De Mots 1965). Samples were mixed thoroughly and filtered (volume of V_S_ sample) through a glass fiber filter. Filter paper was torn into pieces and placed in the extraction vessel; an exact volume V_E_ of 95% ethanol was dispensed into the extraction vessel and the filter pieces were allowed to submerge; the screw cap was closed and resuspended the filter residue by shaking. The vessel was heated in the water bath at 75 ℃ for 5 min, shaken slightly; cooled to room temperature for 15 min; and centrifuged for 10 min at 3000 rpm to obtain a clear supernatant. Measured the absorbance of clear extract in a 1-cm cuvette at 665 nm, 649 nm, and 750 nm against a reference cell filled with ethanol. Equations for determination of chlorophyll a and b in extracts in 95% ethanol:

$\boldsymbol{c}_{\boldsymbol{a}}\mathbf{=13.70}\left( \boldsymbol{A}_{\mathbf{665}}\mathbf{-}\boldsymbol{A}_{\mathbf{750}} \right)\mathbf{-5.76}\left( \boldsymbol{A}_{\mathbf{649}}\mathbf{-}\boldsymbol{A}_{\mathbf{750}} \right)\left( \boldsymbol{mg}\mathbf{/}\boldsymbol{L} \right)$	(Eq. S1)

$$\boldsymbol{c}_{\boldsymbol{b}}\mathbf{=25.80}\left( \boldsymbol{A}_{\mathbf{649}}\mathbf{-}\boldsymbol{A}_{\mathbf{750}} \right)\mathbf{-7.60}\left( \boldsymbol{A}_{\mathbf{665}}\mathbf{-}\boldsymbol{A}_{\mathbf{750}} \right)\left( \boldsymbol{mg}\mathbf{/}\boldsymbol{L} \right)$$

(Eq. S2)

Calculated the amount of pigment per unit volume as follows:

$\boldsymbol{\rho}\mathbf{=}\frac{\boldsymbol{c}\boldsymbol{\times}\boldsymbol{V}_{\boldsymbol{E}}}{\boldsymbol{V}_{\boldsymbol{S}}}$	(Eq. S3)

where $c_{a}$ and $c_{b}$ were the concentrations of chlorophyll a and chlorophyll b in the ethanol, respectively; $A_{665}$, $A_{649}$, and $A_{750}$ were the absorbances at 665 nm, 649 nm, and 750 nm, respectively; and $\rho$ was the concentration of chlorophyll in the initial sample.

**Text S2: Method for EPS analysis**

EPS was extracted by the modified heating method (Li and Yang 2007). Briefly, 20 mL mixed liquor sample was centrifuged at 5000*g* for 15 min. The supernatant was filtered through a 0.45 μm membrane filter, and the filtrate was collected as the loosely bound EPS (LB-EPS). The pellet after centrifugation was resuspended with 0.9% (w/v) NaCl to the original volume. The mixture was heated at 70 °C for 30 min, followed by centrifugation at 10,000g for 20 min under 4 °C. The supernatant was collected and filtered through 0.45 μm membrane filter, and the filtrate was collected as the tightly bound EPS (TB-EPS). The extracted EPS samples were stored at −20 °C prior to analysis. Extracellular [polysaccharides](https://www.sciencedirect.com/topics/earth-and-planetary-sciences/polysaccharide) (PS) were measured by the phenol–sulfuric acid method (DuBois et al. 1956). Protein (PN) concentrations were measured by the modified Lowry method using Pierce™ Modified Lowry Protein Assay Kit (Thermo Fisher).

**Text S3: Analytical methods in details**

Dissolved oxygen (DO) and pH were measured by pH/RDO/DO meter (Thermo Scientific, Orion Star A216). Concentrations of NH_4_^+^, NO_2_^-^, and NO_3_^-^ were measured by ion chromatography (IC) system (DIONEX AQUINO, Thermo Scientific). Total inorganic nitrogen (TIN) was calculated as the sum of NH_4_^+^-N, NO_2_^-^-N, and NO_3_^-^-N. Chemical oxygen demand (COD) was measured based on standard methods (5200B) with Orbeco-Hellige mid-range (0-1500 mg/L) COD kits. TSS, volatile suspended solids (VSS), and sludge volume index (SVI) were measured according to Standard Methods (Rice et al. 2012). The granular size distribution of the mixed liquor biomass was analyzed with Image J.
